# The cell fate controlling CLE40 peptide requires CNGC9 to trigger highly localized Ca^2+^ transients in *Arabidopsis thaliana* root meristems

**DOI:** 10.1101/2020.11.13.381111

**Authors:** Maike Breiden, Vilde Olsson, Karine Gustavo-Pinto, Patrick Schultz, Gregoire Denay, Jenia Schlegel, Petra Dietrich, Melinka A. Butenko, Rüdiger Simon

## Abstract

Communication between plant cells and their biotic environment is largely dependent on the function of plasma membrane localized receptor-like kinases (RLKs). Major players in this communication within root meristems are secreted peptides, including CLAVATA3/EMBRYO SURROUNDING REGION40 (CLE40). In the distal root meristem, CLE40 acts through the receptor like kinase (RLK) ARABIDOPSIS CRINKLY4 (ACR4) and the leucine-rich repeat (LRR) RLK CLAVATA1 (CLV1) to promote cell differentiation. In the proximal meristem, CLE40 signalling requires the LRR receptor-like protein (RLP) CLAVATA2 (CLV2) and the membrane localized pseudokinase CORYNE (CRN), and serves to inhibit cell differentiation. The molecular components that act immediately downstream of the CLE40-activated receptors are not yet known. Here we show that active CLE40 signalling triggers the release of intracellular Ca^2+^ leading to increased cytosolic Ca^2+^ concentration ([Ca^2+^]_cyt_) in a subset of proximal root meristem cells. This rise in [Ca^2+^]_cyt_ depends on the CYCLIC NUCLEOTIDE GATED CHANNEL 9 (CNGC9), CLV1, the CLV1-related BARELY ANY MERISTEM1 (BAM1), CLV2 and CRN. The precise function of changes in [Ca^2+^]_cyt_ are not yet known, but might form a central part of a fine-tuned response to CLE40 peptide that serves to integrate root meristem growth with stem cell fate decisions and initiation of lateral root primordia.

## Introduction

The growth and development of multicellular organisms such as plants is highly coordinated via cell-to-cell interactions. Small secreted peptide ligands, together with different plant hormones, play an important role in maintaining this communication, by fulfilling a vast variety of functions in development, growth and stress responses (Matsubayashi 2011; Olsson et al. 2018). One family of peptide ligands belonging to the class of posttranslationally modified peptides (Matsubayashi 2011; Olsson et al. 2018), is the CLE (CLAVATA3/EMBRYOSURROUNDING REGION, CLV3/ESR) family (Lease and Walker 2006). These peptides perform diverse roles in developmental signalling processes, but also inform on nutrient availability or interactions with beneficial microbes (Cock and McCormick 2001; Gutierrez-Alanis et al. 2017; Mortier et al. 2010; Okamoto et al. 2009).

Here we focus on the role of CLE40, a close homologue of the shoot stem cell secreted peptide CLAVATA3 (CLV3). CLE40 and CLV3 control homeostasis of root or shoot stem cell populations, respectively (Fletcher et al. 1999; Hobe et al. 2003). In the shoot apical meristem (SAM), CLV3 is secreted from stem cells at the meristem tip, and interacts with the plasma membrane localized receptor kinase CLV1, a receptor heteromer consisting of CLV2 and CRN, and the RECEPTOR-LIKE PROTEIN KINASE 2/TOADSTOOL2 (RPK2) (Bleckmann et al. 2010; Fletcher et al.; Jeong et al. 1999; Kinoshita et al. 2010; Müller et al. 2008). Interaction of CLV3 with these receptors ultimately regulates the expression of *WUS* (*WUSCHEL*), encoding a homeodomain transcription factor (TF), which is expressed in the organizing centre (OC) of the SAM (Brand et al. 2000; Mayer et al. 1998; Schoof et al. 2000). The WUS protein was shown to move from its site of expression in the OC to the meristem tip, where it serves to maintain stem cells in an undifferentiated state, and binds to the *CLV3* promoter to increase *CLV3* expression levels (Daum et al. 2014; Yadav et al. 2011). The negative regulation of *WUS* expression by CLV3 and positive regulation of *CLV3* expression by WUS establishes a negative feedback loop that sustains stem cell homeostasis (Brand et al. 2000; Schoof et al. 2000).

Another target of regulation by CLV3 is expression of the CLV1-related RLK BAM1. The extracellular LRRs of BAM1 can, similar to those of CLV1, recognize and bind CLV3 peptide (Shinohara and Matsubayashi 2015). Under normal conditions, *BAM1* is expressed at the flanks of the SAM with little or no expression in the centre of the meristem where *CLV1* is expressed (DeYoung et al. 2006). In *clv1* mutant meristems, *BAM1* becomes expressed in a domain comprising the OC, where BAM1 can partially compensate for loss of CLV1 to restrict *WUS* expression (Nimchuk et al. 2015). If both *BAM1* and *CLV1* are mutated, stem cells accumulate resulting in uncoordinated growth of the meristem (DeYoung and Clark 2008; Nimchuk et al. 2015). The precise role of the other receptor systems comprising CLV2/CRN and RPK2 is less clear. More stem cells and higher *WUS* expression were found in *clv2*, *crn* or *rpk2* mutants, and various studies reported that larger receptor protein complexes between CLV1, CLV2/CRN, BAM1 and RPK2 can form at membranes (Betsuyaku et al. 2011; Kayes and Clark 1998; Kinoshita et al. 2010; Bleckmann et al., 2010; Müller et al. 2008; Shimizu et al. 2015; Somssich et al. 2015). Downstream of these receptor complexes, heterotrimeric G-proteins and MAP-kinases were found to contribute to further signal processing and transmission (Betsuyaku et al. 2011; Bommert et al. 2013; Ishida et al. 2014).

The fate of stem cell population in the distal root meristem depends on the presence of CLE40 in differentiated columella cells (CCs) negatively regulating the number of columella stem cells (CSCs) via ACR4 and CLV1 in a quantitative manner (Stahl et al. 2009; Stahl et al. 2013). In the proximal root meristem, CLE40 is required in the stele to inhibit or delay meristem cell differentiation. CLE40 induced signalling affects the expression of several phytohormone biosynthetic genes and stem cell specific transcription factor genes such as *WUSCHEL-LIKE HOMEOBOX5* (*WOX5)*, but the components of the intracellular signalling cascade triggered by CLE40 are so far unknown. Curiously, increased *CLE40* expression or external addition of CLE40 peptide (CLE40p) also causes premature meristem differentiation (like the loss-of-function situation), indicating that root meristem maintenance might strongly depend on CLE40 concentration (Hobe et al. 2003; Pallakies and Simon, 2014; Stahl et al. 2009).

CLE family peptides act in both local and systemic communication pathways and share central components with pathogen response pathways, such as small secreted ligands, plasma membrane localized RLK complexes and downstream acting cytoplasmic kinases or G-proteins that ultimately converge on regulation of target gene expression (Olsson et al. 2018). Pathogen responses often involve the generation of reactive oxygen species (ROS). ROS can serve as a first line of defence, and ROS production is intimately linked to an increase in cytosolic Ca^2+^ concentration ([Ca^2+^]_cyt_) (Demidchik et al. 2003; Pei et al. 2000). In resting cells [Ca^2+^]_cyt_ are kept at a steady state of around 100 nM, while the apoplast maintains a 1 mM Ca^2+^ concentration (Stael et al. 2012). Upon diverse developmental or environmental signals such as drought stress, high salt or auxin, the [Ca^2+^]_cyt_ can rise rapidly and activate a large toolset of Ca^2+^ sensor proteins that affect protein phosphorylation or gene expression (Reviewed in (Kudla et al. 2018)). Ca^2+^ influx across the plasma membrane is mediated by different families of Ca^2+^ permeated channels including CYCLIC NUCLEOTIDE GATED CHANNELs (CNGCs) and GLUTAMATE LIKE RECEPTORs (GLRs) (Clough et al. 2000).

Ca^2+^ influx can be initiated by activation of plasma membrane RLKs, for example, interaction of a flagellin peptide (flg22) with the extracellular domain of the LRR-RLK FLAGELLIN SENSING2 (FLS2) leads to an increase in the [Ca^2+^]_cyt_ and activation of downstream components (Boller and Felix 2009; Chinchilla et al. 2007). BOTRYTIS-INDUCED KINASE1 (BIK1), a receptor-like cytoplasmatic kinase that interacts with FLS2 and the co-receptor BRASSINOSTEROID INSENSITIVE1-ASSOCIATED RECEPTOR KINASE1/SOMATIC EMBRYOGENESIS-RELATED KINASE3 (BAK1/SERK3), is rapidly released from the plasma membrane complex upon flg22 perception and activates the NADPH oxidase RESPIRATORY BURST OXIDASE PROTEIND (RBOHD) by phosphorylating its N-terminus (Kadota et al. 2014; Li et al. 2014). Activated RBOHD catalyses a Ca^2+^ influx via the production of ROS, which also further aids in regulating the activity of RBOHD via CALCIUM DEPENDENT PROTEIN KINASE5 (Dubiella et al. 2013). In addition, there can be microbe-or damage-associated molecular patterns triggered increase in [Ca^2+^]_cyt_ independent of the presence of functional RBOHD. However, the Ca^2+^ response is attenuated compared to when RBOHD is present, demonstrating a feedback effect of ROS on Ca^2+^ signalling (Ranf et al. 2011).

flg22 induced Ca^2+^ efflux from the cytoplasm is mediated via the AUTOINHIBITED Ca^2+^ ATPase8 (ACA8) and ACA10 which were shown to interact with FLS2 after flg22 binding (Frei dit Frey et al. 2012). Ca^2+^ activated proteins mostly contain an EF hand Ca^2+^ binding domain (Day et al. 2002; Kudla et al. 2018), and are activated by different Ca^2+^ concentrations due to their specific affinities (Geiger et al. 2010; Scherzer et al. 2012).

The close interplay between ROS production and intracellular Ca^2+^ release can serve signal amplification and rapid, systemic signal propagation from a single perceiving cell to distal sites and organs (Steinhorst and Kudla 2013). Both ROS production and Ca^2+^ release were recently shown to mediate downstream signalling of the INFLORESCENCE DEFICIENT IN ABSCISSION (IDA) peptide, which is closely related to CLE peptides and acts at the interface between developmental and pathogen response pathways by coordinating cell separation with immune responses (Olsson et al. 2019). Although the intracellular events from plasma membrane receptor activation to final cellular output leading to stem cell maintenance is far from understood, a recent study indicated that Ca^2+^ level fluctuations influence auxin distribution and PIN-FORMED1 (PIN1) polarity in the SAM of Arabidopsis (Li et al. 2019). In addition, Ca^2+^ signalling has been suggested to act as a second messenger in CLV signalling in Arabidopsis seedlings (Chou et al. 2016).

In this study we focused on analysing whether CLE40 can induce elevations in [Ca^2+^]_cyt_ using genetically encoded Ca^2+^ indicators (GECIs) which are best suited for *in vivo* Ca^2+^ measurements. Numerous GECIs have been developed since the discovery of aequorin (Shimomura et al. 1962) and first design of Cameleons (Miyawaki et al. 1997) that can be categorized into single fluorescent protein (FP) intensiometric and two-FP ratiometric sensors (Pérez Koldenkova and Nagai 2013). The sensor used in this study, R-GECO1, is a red-emitting single FP (Zhao et al. 2011) and has been shown to exhibit an increased dynamic range compared to another commonly used sensor, the FRET-based Cameleon YC3.6 (Keinath et al. 2015).

We here investigated whether Ca^2+^ is a component of CLE40 signalling. We demonstrate that CLE40 signalling leads to a rapid, but spatially highly localized increase in [Ca^2+^]_cyt_ in a subgroup of Arabidopsis root meristem cells. Using mutants and inhibitor studies, we show that CLV1 and BAM1 are both required for CLE40 dependent Ca^2+^ release, while the CLV2/CRN heteromer contributes to a lesser extent. We also show that CNGC9 is one of the key channels that are activated by CLE40 signalling in root cells and that loss of CNGC9 activity interrupts Ca^2+^ transients while root meristem differentiation remains unaffected.

## Results

### CLE40 peptide induces a distinct, localized increase in intracellular Ca^2+^ concentration

Root meristems respond in a highly sensitive manner to changes in the concentration of CLE40 peptide (CLE40p) (Fiers et al. 2005; Pallakies and Simon 2014), indicating that seedlings respond to exogenous application of CLE40p by biological relevant readouts. Treatment of Col-0 wild type (WT) seedlings with 200 nM of CLE40p leads to shorter roots due to premature differentiation of the proximal root meristem compared to untreated WT roots (Supplementary Fig.1 A,B) (Hobe et al. 2003).

To investigate whether CLE40 signalling involves changes in [Ca^2+^]_cyt_, we used Arabidopsis plants expressing R-GECO1 as a Ca^2+^ sensor (Keinath et al. 2015). By continuously monitoring primary roots at 10 days after germination (DAG) in an imaging chamber via an inverted confocal microscope we could detect an increase in the normalized fluorescence intensity of R-GECO1 in a small group of cells in the proximal root meristem upon addition of 1 µM CLE40p (Movie 1 and Fig.1 A,B). The CLE40p induced Ca^2+^ signals become visible in the stele region of the root within 7 minutes (min) after addition of CLE40p to the chamber. Here, small regions responded to CLE40p with rapid spikes in their fluorescence intensity that was quantified by placing regions of interest (ROIs) in the stele (Movie1 and Fig.1 A,B). We have so far not able to resolve if these signals originate from individual cells, or small cell groups that respond simultaneously. This local [Ca^2+^]_cyt_ elevation was noted in about 55 % of all roots tested (n = 62), but missing in 45 % of all roots that still responded to control extracellular ATP (eATP) treatment (see below) (Table 1). The frequency of roots responding to CLE40p is in a similar range as that observed for chitin treated roots, where 67 % of roots showed [Ca^2+^]_cyt_ elevations (Keinath et al. 2015).

**Fig. 1:**
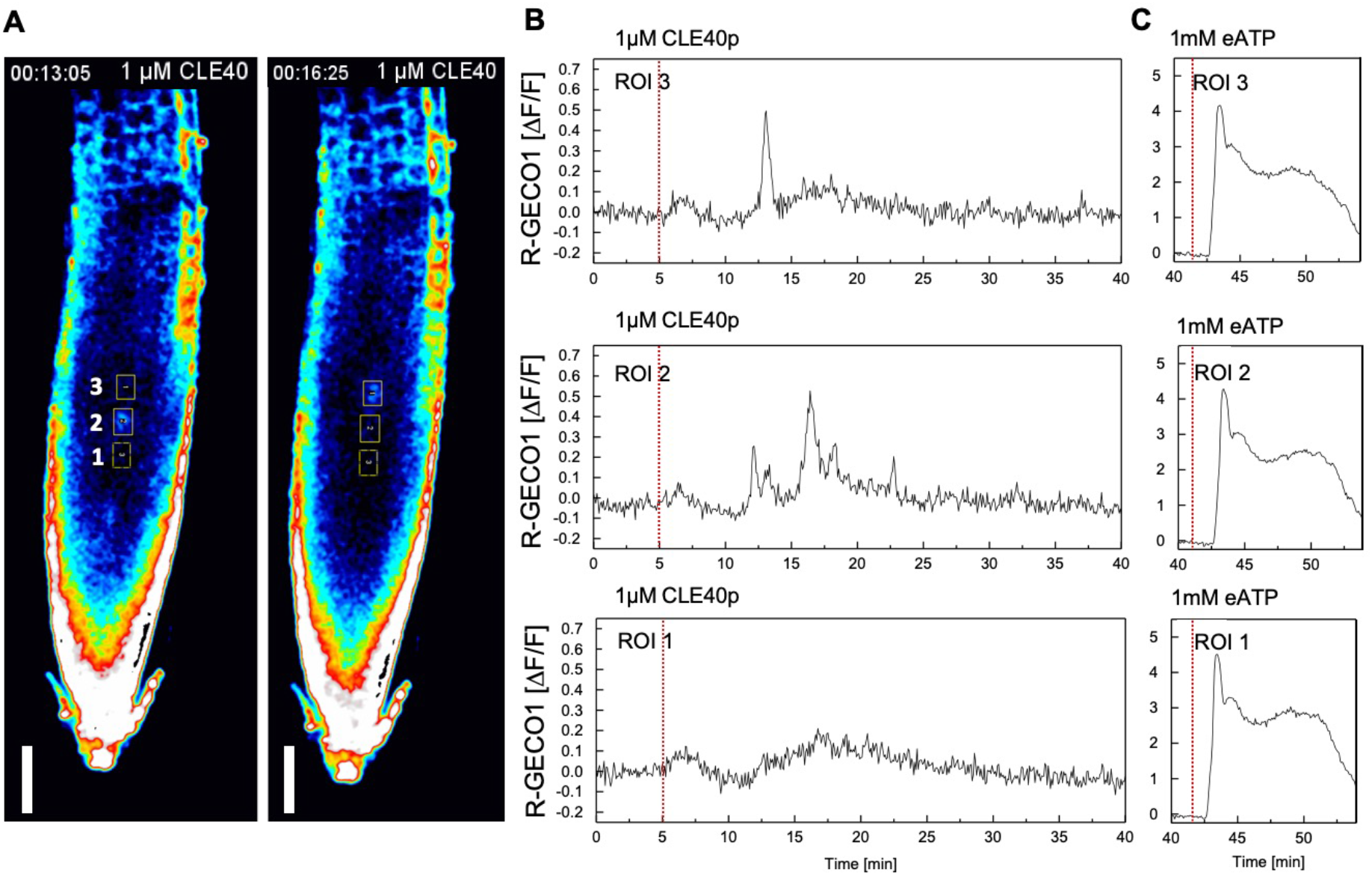
CLE40p treatment induces a distinct increase in [Ca^2+^]_cyt_. **A**, Representative response of the changes in [Ca^2+^]_cyt_ observed when roots expressing *R-GECO1* are treated with 1 µM CLE40p. The increase in [Ca^2+^]_cyt_ response is observed as spikes in a small group of cells in the stele region. ROIs (outlined in yellow rectangles) were used to measure normalized R-GECO1 fluorescence intensities (∆F/F). Cells with increase in [Ca^2+^]_cyt_ in ROI 3 and ROI 2 are observed at time point 13:05 (mm:ss) and 16:25 (mm:ss), respectively. No changes in [Ca^2+^]_cyt_ are observed in ROI 1. Scale bar = 50 µm. **B**, Normalized R-GECO1 fluorescence intensities (∆F/F) were measured from ROI 1, ROI 2 and ROI 3 in the meristematic zone of the root. Shown are ([Ca^2+^]_cyt_) dynamics in response to 1 µM CLE40p over time (see also Movie 1). Red line at 5 min indicates application of CLE40p. **C**, Normalized R-GECO1 fluorescence intensities (∆F/F) were measured from ROI 1, ROI 2 and ROI 3 in the meristematic zone of the root. Shown are [Ca^2+^]_cyt_ dynamics in response to 1 mM eATP over time (see also Movie 1). The addition of 1 mM eATP serves as an internal positive control and leads to a rapid increase in [Ca^2+^]_cyt_ and increased fluorescence intensity. **A,B,C,**representative response from 62 roots. Red line at 5 min indicates application of CLE40p or at 42 min application of eATP, respectively.

**Table 1.** CLE40p and CLE40p-R induced [Ca^2+^]_cyt_ increase in Col-0 WT and receptor mutant lines.

| Line | n | [Ca<sup>2+</sup>]<sub>cyt</sub> Responses (n) |  |  | [Ca<sup>2+</sup>]<sub>cyt</sub> Responses – % |  |  |
| --- | --- | --- | --- | --- | --- | --- | --- |
|  |  | yes | unclear | no | yes | unclear | no |
|  |  | CLE40p Treatment |  |  |  |  |  |
| Col-0 | 62 | 34 | 7 | 21 | 55 | 11 | 34 |
| *bam1-3* | 13 | 2 | 0 | 11 | 15 | 0 | 85 |
| *clv1-20* | 10 | 1 | 1 | 8 | 10 | 10 | 80 |
| *clv2-gabi* | 14 | 5 | 0 | 9 | 36 | 0 | 64 |
| *crn-3* | 14 | 3 | 0 | 11 | 21 | 0 | 79 |
|  |  | CLE40p-R Treatment |  |  |  |  |  |
| Col-0 | 14 | 2 | 1 | 11 | 14 | 7 | 79 |

Addition of a randomized CLE40 peptide (CLE40p-R) with the same amino acid composition as CLE40p but a different amino acid sequence, to the imaging chamber triggered no changes in [Ca^2+^]_cyt_ even if the roots responded to control eATP treatment (see below), indicating that an active CLE40p was essential for the [Ca^2+^]_cyt_ elevation (Table 1).

Since [Ca^2+^]_cyt_ transients can be triggered by diverse biotic and abiotic stresses including mechanical stresses and other endogenous peptides, we wanted to assay for the specificity of the CLE40p responses. To investigate whether [Ca^2+^]_cyt_ spikes are limited to treatment with CLE40p, we used a closely related peptide, CLE14 peptide (CLE14p). CLE14p is expressed in the root cap of Arabidopsis and involved in signalling of phosphate deficiency (Gutierrez-Alanis et al. 2017). Upon phosphate starvation, *CLE14* expression is upregulated and expands to the endodermis and stele. Given the localized [Ca^2+^]_cyt_ elevation to a small number of cells after CLE40p treatment (Movie 1 and Fig.1 A,B) we reasoned that we needed to increase the ROIs along the whole stele to compare signal intensity values between different treatments. When comparing signal intensity values from roots treated with 1 µM CLE40p to 1 µM CLE14p in ROI s along the stele we could detect increased [Ca^2+^]_cyt_ after application of CLE40p, but not after application of CLE14p (Fig.2 A,B), indicating that eliciting a [Ca^2+^]_cyt_ elevation is not a general feature of CLE peptide signalling pathways. We noted that roots at 10 DAG were more responsive to CLE40p than younger roots, and therefore suggest that a key component in the CLE40 signalling pathway might be rate limiting and be only lowly expressed at earlier stages.

**Fig. 2:**
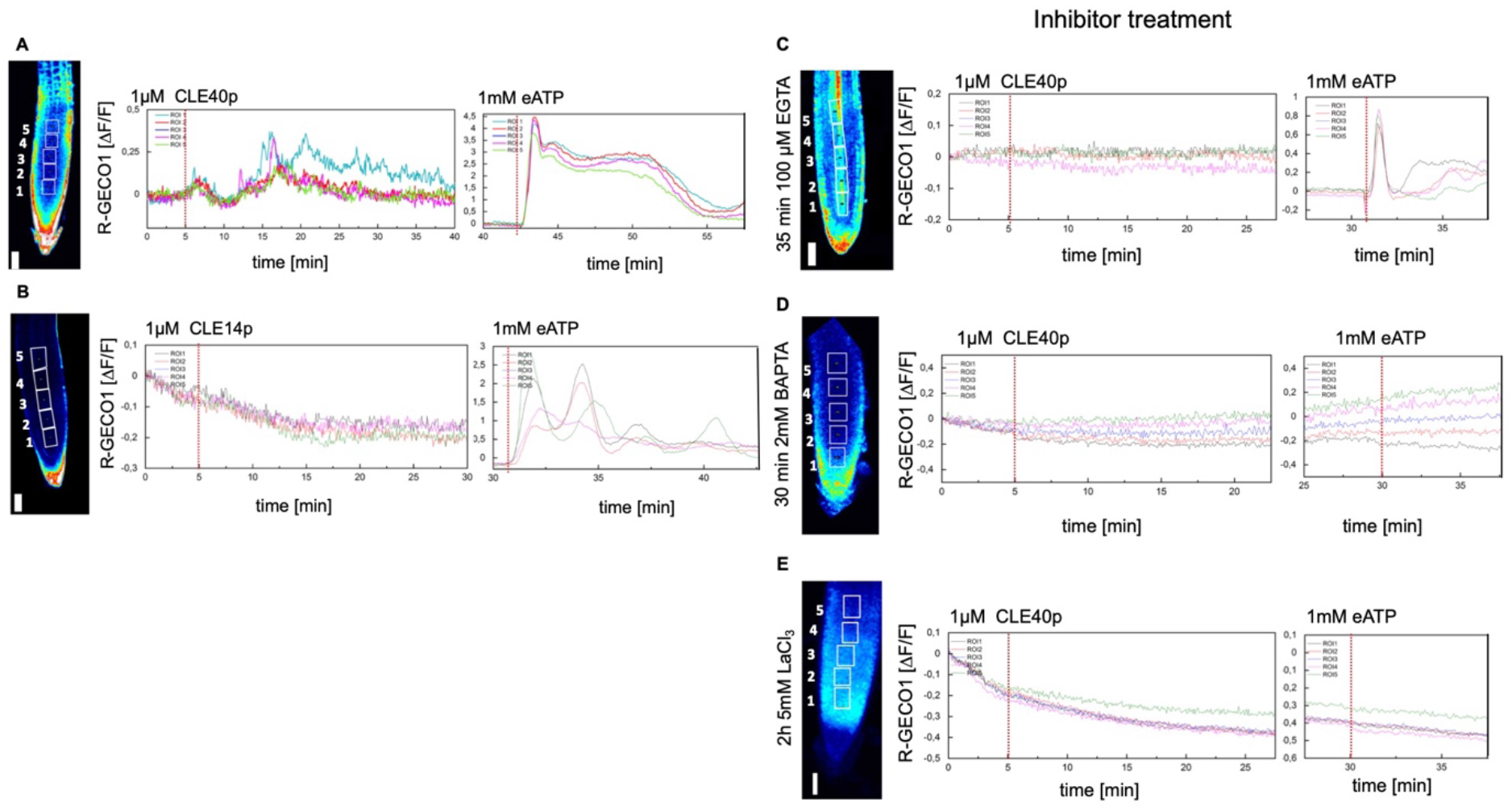
CLE40p induced increase in [Ca^2+^]_cyt_ can be blocked by inhibitors. Representative response of the changes in [Ca^2+^]_cyt_ observed when roots expressing *R-GECO1* are treated with 1 µM CLE40p or 1 µM CLE14p followed by 1mM eATP. ROI 1-5 (outlined in white rectangles) were used to measure normalized R-GECO1 fluorescence intensities (∆F/F) and are represented in the graphs as dynamic responses over time. Red line at 5 min indicates application of CLE40p, followed by an additional red line indicating application of eATP, used as an internal positive control. **A**, Increase in [Ca^2+^]_cyt_ response is observed as spikes after treatment with CLE40p. Cells with increase in [Ca^2+^]_cyt_ in ROI 4 are observed at time point 12 min, followed by additional spikes in ROIs 1-5. Same root as depicted in Fig. 1 A,B but with larger ROI s analysed, n = 62. **B**, No change in [Ca^2+^]_cyt_ response is observed in ROI 1-5 after treatment with CLE14p, n = 17. **C**, No change in [Ca^2+^]_cyt_ response is observed in ROI 1-5 when roots were pre-incubated for 30 min in 100 µM EGTA, a Ca^2+^ chelator, before treatment with CLE40p, n = 14 **D**, No change in [Ca^2+^]_cyt_ response is observed in ROI 1-5 when roots were pre-incubated for 60 min in 2 mM BAPTA, an intracellular Ca^2+^ chelator, before treatment with CLE40p, n = 11. **E**, No change in [Ca^2+^]_cyt_ response is observed in ROI 1-5 when roots were pre-incubated for 120 min in 5 mM LaCl_3_, blocking plasma membrane localized Ca^2+^ channels, before treatment with CLE40p, n = 10. Addition of 1mM eATP leads to increased fluorescence intensity representing an increase in [Ca^2+^]_cyt_ for **B**and **C**, while there is no increased fluorescence intensity upon eATP treatment in **D and E**. **(A, B, C, D, E)** Scale bar = 50 µM.

### The CLE40p triggered [Ca^2+^]cyt response requires Ca^2+^ influx from the apoplast

Further dissection of the CLE40 induced [Ca^2+^]_cyt_ elevation required an independent positive control that triggers a Ca^2+^ response. The flg22 peptide which acts via the FLS2 receptor and the BAK1 co-receptor has already been shown to cause a strong Ca^2+^ signal, which moves as a response wave through the entire root tip, starting from the root elongation zone and further spreading towards the meristematic zone (Keinath et al. 2015). However, the intensive cross-talk between PAMP-triggered and developmental signalling pathway, exemplified by the activation of common target genes by flg22 and the IDA peptide, which also involves similar ROS and Ca^2+^ responses (Olsson et al. 2019), can compromise further analysis of downstream signalling events. Osmotic- and salt-stresses stimulate Ca^2+^ release, but NaCl leads to a rapid shrinkage of the root meristem due to the change in water potential, confounding image analysis (Knight et al. 1997; Munns and Passioura 1984). We therefore used eATP as a reference stimulus for [Ca^2+^]_cyt_ release. eATP serves as a danger signal and is sensed via the plasma membrane localized receptor-kinase DOES NOT RESPOND TO NUCLEOTIDES1 (DORN1/P2K1) (Choi et al. 2014; Keinath et al. 2015; Waadt et al. 2017). Adding 1 mM eATP caused reproducible [Ca^2+^]_cyt_ elevation within 1 min that initiated in the root tip and extended shootwards as a wave (Fig.1 C, Fig.2 A-E and Fig.3). In contrast to CLE40p induced [Ca^2+^]_cyt_ elevation, treatments with eATP, flg22, and IDA affect the entire root meristem and were not locally restricted. In addition, the gap period between stimulus and [Ca^2+^]_cyt_ response varies considerably between treatments, i.e., 7 min for CLE40p peptide and 1 min for eATP (Fig.1 C, Fig.2 A-E and Fig.3), which indicates mechanistic differences between the response systems involved. eATP triggered [Ca^2+^]_cyt_ elevation in all cells including the epidermis, while CLE40p elicited a response only in small regions within the stele. The longer diffusion time required for CLE40p to reach target sites deeper in the root tissue could be one explanation for this difference; the utilization of different signalling components for Ca^2+^ perception and transmission between different stimuli could be another.

**Fig. 3:**
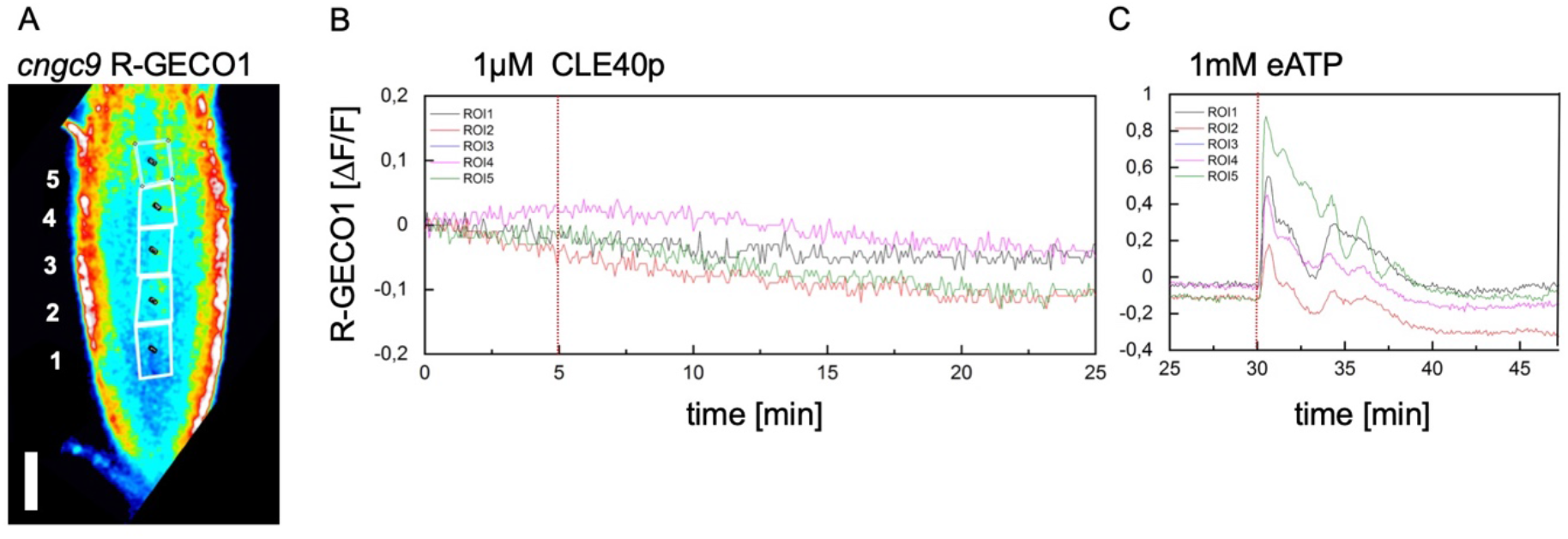
CNGC9 is necessary for the CLE40p induced [Ca^2+^]_cyt_ increase. Representative response of the changes in [Ca^2+^]_cyt_ observed when roots expressing *R-GECO1* in the *cncg9* mutant background are treated with 1 µM CLE40p followed by 1 mM eATP. **A**, ROI 1-5 (outlined in white rectangles) were used to measure normalized R-GECO1 fluorescence intensities (∆F/F) and are represented in the graphs as dynamic responses over time. **B**, Red line at 5 min indicates application of CLE40p. **C**, red line at 30 min indicates application of eATP, used as an internal positive control. No change in [Ca^2+^]_cyt_ response is observed in ROI 1-5 after treatment with CLE40p, while addition of eATP leads to increased fluorescence intensity representing an increase in [Ca^2+^]_cyt_, n = 9. Scale bar = 50 µm.

We tested three Ca^2+^ signalling inhibitors to further investigate if the CLE40p induced signal was generated by Ca^2+^ influx from the extracellular space. LaCl_3_ is able to block cation channels in the plasma membrane and thereby inhibits an [Ca^2+^]_cyt_ elevation (Knight et al. 1997), for example in response to eATP (Wang et al. 2014). Ethylene glycol-bis(2-aminoethylether)-N,N,N′,N′-tetraacetic acid (EGTA) is able to chelate Ca^2+^ that is present in the extracellular space (Knight et al. 1997). As a third inhibitor 1,2-bis(o-aminophenoxy)ethane-N,N,N’,N’-tetraacetic acid (BAPTA), a ph-insensitive Ca^2+^ chelator (Tsien 1980, Li et al. 2019) was used. Pre-incubation of roots with either LaCl_3,_ EGTA or BAPTA inhibited CLE40p induced [Ca^2+^]_cyt_ elevation (Fig.2 C, D, E). Pre-incubation in 100 µM EGTA gave no response of the roots to CLE40p, however, eATP still triggered a minor rise in [Ca^2+^]_cyt_, which shows that the small and very localized [Ca^2+^]_cyt_ elevation stimulated by CLE40p is highly sensitive to a reduction in free extracellular Ca^2+^ levels (Fig.2 C). Preincubation with BAPTA (Fig. 2 D) as well as pre-incubation with LaCl_3_ (Fig.2 E) lead in all cases to no visible response, neither to CLE40p or eATP addition.

### CNGC9 is required for CLE40p induced elevation of [Ca^2+^]_cyt_

The lack of [Ca^2+^]_cyt_ response in LaCl_3_ treated roots indicated that the CLE40p dependent [Ca^2+^]_cyt_ response required plasma membrane localized Ca^2+^ channel. In order to identify the Ca^2+^ channel responsible for the CLE40p triggered increase in [Ca^2+^]_cyt_, we analysed mutants for all root-expressed CNGCs in their reaction to CLE40p. Mutant lines grown on agar containing 50 nM CLE40p responded with a premature differentiation of meristem cells and reduced root growth. The reaction of mutant roots was similar to that of WT, indicating that none of the CNGCs is required for this response (Supplementary Fig. 3).

We then tested the role of individual CNGCs in the [Ca^2+^]_cyt_ elevation. We found that a transcriptional null mutant of *CNGC9* (Brost et al. 2019), carrying the R-GECO1 reporter, did not show a Ca^2+^ release in response to CLE40p, while the response to eATP was still detectable (Fig.3). We conclude that CNGC9 is an essential channel that opens in response to CLE40p signalling and permits rapid Ca^2+^ influx.

### CLE40 signals via distinct membrane receptors to control [Ca^2+^]_cyt_ or stem cell fate

We previously noted that CLE40 acts through several different receptors in distinct cell fate control pathways. In the distal root meristem, CLE40 controls the differentiation status of CSCs via CLV1 and ACR4 (Stahl et al. 2009; Stahl et al. 2013), while regulation of *WOX5* expression proximal to the quiescent centre (QC) depends on the CLV2/CRN heteromer (Berckmans et al. 2019). Given that CLE40 signalling in the proximal root meristem requires BAM1, CLV1, CLV2 and CRN we investigated the expression pattern of these receptors by promoter activity or promoter-protein fusion expression. The promoters of *CLV2* and *CRN* were cloned and fused to the nuclear localized *YFP-*derived fluorophore Venus protein or the mCherry protein; while the promoter and coding region of *BAM1* and *CLV1* were fused to GFP, respectively. To analyse the expression pattern of *pBAM1:BAM1-GFP*, *pCLV1:CLV1-GFP*, *pCLV2: H2B*-*Venus* and *pCRN: H2B-mCherry*, roots were stained with propidium iodide and investigated using a confocal laser scanning microscope. BAM1 expression was found in the proximal meristem, while CLV1 was expressed in the CSC of the distal meristem (Fig.4 A,B). CLV1 expression is also detected in the phloem companion cells in the stele (Araya et al. 2014). Promoter activity was detected for *CLV2* and *CRN* in cells of the proximal meristem and the promoter of *CLV2* had prominent activity in the stele (Fig.4 C,D). Interestingly, when comparing the expression pattern of *BAM1*, *CLV1*, *CLV2* and *CRN* with the expression of *CLE40* from roots expressing a *pCLE40:H2B-Venus* construct we observed that the receptor expression overlaps with those of *CLE40* in the proximal root meristem and the stele where the CLE40p induced increase in [Ca^2+^]_cyt_ is visible (Fig.4 A-E).

**Fig. 4:**
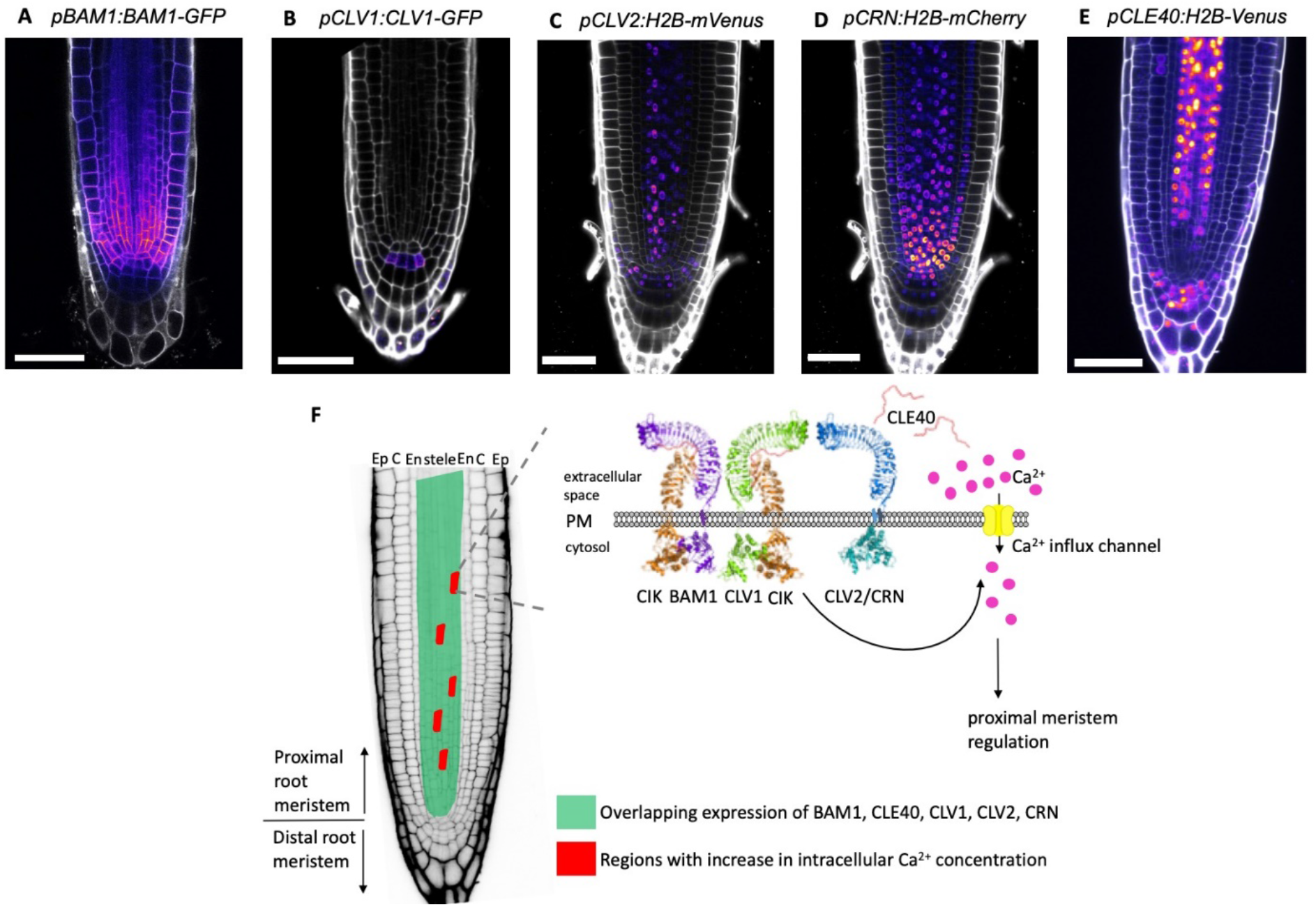
Expression patterns of *BAM1, CLV1, CLV2, CRN* and *CLE40* in the root meristem. Microscopic analysis of 2 - 10 dag *pBAM1:BAM1-GFP*, *pCLV1:CLV1-GFP*, *pCLV2:H2B-mVenus*, *pCRN:H2B-mCherry* and *pCLE40:H2B-Venus* roots. **A**, *pBAM1:BAM1-GFP* expressing roots show expression of the BAM1 receptor in the proximal meristem, 2 dag. **B**, *pCLV1:CLV1-GFP* expressing roots show expression of the CLV1 receptor in the columella stem cells (CSCs), 10 dag. **C**, Fluorescent nuclei could be observed in the stele of the proximal meristem, in the CSCs and columella cells (CCs) in *pCLV2:H2B-mVenus* expressing roots, 5 dag. **D**, Fluorescent nuclei could be observed in the all cell layers of the proximal meristem except for the epidermis in *pCRN:H2B-mCherry* expressing roots, 5 dag. **E**, Fluorescent nuclei could be observed in the stele of the proximal meristem and CCs in *pCLE40:H2B-Venus* expressing roots, 10 dag. For **C** and **D** roots co-expressing *pCLV2:H2B-mVenus* and *pCRN:H2B-mCherry* were used. **A,B,C,D,**Scale bars = 50 µm. Cell walls of the roots were counterstained using propidium iodide and are depicted in grey. Fluorescence of reporter lines are depicted using the look up table fire. Maximum intensity projections of z-stacks**. E,**Model for how CLE40 induces increase in [Ca^2+^]_cyt_ in the proximal meristem. The increase in [Ca^2+^]_cyt_ can be seen in a subset of cells in the stele region (highlighted in red) of the root where *BAM1, CLV1, CLV2*, *CRN* and *CLE40* are expressed. The activation of the receptors, in the presence of the CIK (CLAVATA3 INSENSITIVE RECEPTOR KINASE) coreceptors (Hu et al. 2018), upon CLE40 binding or interaction leads to the activation of the CNGC9 Ca^2+^ influx channel in the plasma membrane. The influx of Ca^2+^ ions then acts via a still unknown signalling cascade to regulate the proximal root meristem. Changes in [Ca^2+^]_cyt_ are not observed in the distal root meristem where the CLE40 signal is transmitted via the receptors CLV1 and ACR4 to regulate the distal root meristem.

To further explore the role of these receptors in transmitting the CLE40 signal leading to an increase in [Ca^2+^]_cyt_ we crossed the R-GECO1 reporter into receptor mutant lines and assayed their response to the addition of CLE40p. *acr4* mutants were not included in this study since *ACR4* expression in the proximal root meristem is predominantly epidermal (Gifford et al. 2003), where no [Ca^2+^]_cyt_ elevation was noted. The CLE40p induced increase in [Ca^2+^]_cyt_ in the stele region of the seedlings is reduced in all receptor mutant backgrounds, compared to WT plants (Table 1). In *bam1-3* and *clv1-20* background, only 10 % or 15 %, respectively, of all tested roots responded, while 55 % of all WT roots displayed a rapid rise in [Ca^2+^]_cyt_ (Table 1 and Supplementary Fig.2 A,B). The requirement for the CLV2/CRN heteromer was less pronounced, and 21 % to 36 % of all roots still responded to CLE40p in the *clv2-gabi* and *crn3* mutant background, respectively (Table 1 and Supplementary Fig.2 C,D). Thus, none of the mutants in individual receptors fully abolished the Ca^2+^ response to CLE40p. A possible interpretation is that CLE40p acts via several receptors, and that single mutants therefore still allow for some signalling through the Ca^2+^ mediated pathway. Alternatively, mutual transcriptional regulation could compensate for the loss of receptors, as was previously described for the upregulation of *BAM1* expression in *clv1* mutants in the SAM (Nimchuck et al. 2015).

## Discussion

CLE40 signalling in the root meristem involves the receptors ACR4 and CLV1 in the distal root meristem and BAM1, CLV1, CLV2 and CRN in the proximal root meristem (Fig.4 F) (Pallakies and Simon 2014; Stahl et al. 2009; Stahl et al. 2013). CLE40 perception leads to the promotion of cell differentiation and loss of CSCs in the distal root meristem (Stahl et al. 2009). In the proximal root meristem, CLE40 inhibits cell differentiation (Pallakies and Simon 2014). Downstream signalling components of the CLE40 signalling cascade are not yet known. Using a red-emitting single-fluorescent genetically encoded Ca^2+^ indicator (R-GECO1), we here show that the perception of CLE40p leads to an increase in [Ca^2+^]_cyt_. Ca^2+^ signals were observed in the stele region of the root meristem, and spatially confined to a small number of cells (Fig.1 A,B). The increase in [Ca^2+^]_cyt_ can be blocked using the inhibitors LaCl_3_, EGTA and BAPTA (Fig.2 C,D,E). Ca^2+^ influx is therefore most likely starting from the apoplastic space and mediated via Ca^2+^ permeable channels in the plasma membrane. Ca^2+^ influx from the apoplast was specifically triggered by CLE40p, and not observed upon the addition of other root active CLE peptides, such as CLE14p (Fig.2 B). Furthermore, we found that the CLE40 receptors which are required to promote root meristem differentiation in response to an increased dosage of CLE40p (Pallakies and Simon 2014; Stahl et al. 2013) were also required for the rise in [Ca^2+^]_cyt_, indicating that the changes in cell states and Ca^2+^ signalling are intimately linked (Table 1 and Supplementary Fig.2 A-D).

Our screen for CNGC proteins that might control [Ca^2+^]_cyt_ resulted in the identification of CNGC9 as a key channel protein (Fig.3). An emerging scenario is that CLE40p activation of membrane localized RLK complexes comprising CLV1 and the CLV2/CRN heteromer results in rapid phosphorylation of CNGC9 and, in consequence, elevation of [Ca^2+^]_cyt._ The observation that only a very small number of cells respond to the CLE40p signal is intriguing, because the *RLKs* and *CNGC9* are expressed in widely overlapping domains in the stele (Fig. 4 A-E) (Brost et al. 2019; Klepikova et al. 2016). We have to postulate that other, highly localized factors restrict the [Ca^2+^]_cyt_ response. We did not test the impact of CIK on the CLE40p induced elevation in [Ca^2+^]_cyt_ since this would have required a quadruple mutant line of *CIK1,2,3,4*. Whether the cells reacting with [Ca^2+^]_cyt_ elevation are the precursor of lateral root primordia is an intriguing hypothesis, but requires further investigation. In our model we propose that the CLE40p triggered increase in [Ca^2+^]_cyt_ is dependent on receptors known to regulate the proximal root meristem as well as CNGC9 (Fig. 4 F), and it will be of interest to investigate whether any of these signalling components are affected in lateral root development.

It was previously shown that CLV3, a close homologue of CLE40, can induce a transient increase in [Ca^2+^]_cyt_ in whole seedlings, which was monitored using the aequorin system for Ca^2+^ detection (Chou et al. 2016). These authors also showed that the induction of [Ca^2+^]_cyt_ is impaired in lines mutant for *CNGC2*, a cation influx channel in the plasma membrane, and that *cngc2* (*dnd1*) mutants are affected in shoot meristem development. Chou and colleagues concluded from their data that [Ca^2+^]_cyt_ elevation is essential for stem cell regulation in the shoot meristem. However, *cngc2/dnd1* mutants are severely retarded in growth and development from embryogenesis onwards, and the failure to maintain a functional meristem cannot be unequivocally assigned to reduced Ca^2+^ signalling (Chou et al. 2016). In our studies, *cngc2* mutants, when grown on growth medium in sterile culture, were indistinguishable from the WT and were not affected in stem cell behaviour (Supplementary Fig.3).

Li et al. (2019) addressed the role of Ca^2+^ signalling in the SAM of *Arabidopsis*. They employed the R-GECO1 and GCaMP6f Ca^2+^ sensors and observed continuous spiking signals in individual shoot meristem cells. The relevance of these signals is still unclear, however, inhibiting Ca^2+^ signalling using a high-dosage treatment with LaCl_3_ or BAPTA affected the repolarization behaviour of the PIN1 auxin efflux carrier upon wounding. The authors concluded that Ca^2+^ spiking could be a prerequisite for the correct polarization of PIN1 protein and may contribute to the establishment of highly ordered phyllotactic patterns in the shoot meristem (Li et al. 2019).

A single EF-hand Ca^2+^-binding protein CA^2+^-DEPENDENT MODULATOR OF ICR1 (CMI1) was identified by Hazak and colleagues in 2019 that is able to regulate auxin responses and affects auxin-induced changes in the cytoplasmic Ca^2+^ levels (Hazak et al. 2019). Auxin up-regulates the expression of CMI1 and CMI1 binds in Ca^2+^-dependent manner to ICR1 (INTERACTOR OF CONSTITUTIVELY ACTIVE ROP). Ca^2+^-dependent binding of ICR1 and CMI1 affects the localization and function of CMI1 leading to recruitment of CMI1 to microtubules in the presence of ICR1. Since CMI1 responds to low Ca^2+^ concentration (<10^-7^ M), and its expression is directly up-regulated by auxin and the loss of function of CMI1 leads to phenotypes associated with impaired auxin responses (repression of auxin-induced Ca^2+^ increases in the vasculature and lateral root cap), CMI1 can therefore rapidly transduce cell-specific auxin signals.

The role of elevations in [Ca^2+^]_cyt_ in CLE peptide signalling is not yet known. The observations reported here point at a localized role in the control of cell identities and fates.

## Material and methods

### Plant material

Plant lines used in this work: Ecotype Colombia-0 (Col-0) was used as wild type (WT). Mutant lines: *clv2-gabi* GK-686A09 and the *cngc* lines were obtained from the Nottingham Arabidopsis Stock Centre (NASC) (*cngc1* SAIL_443_B11, *cngc2* SALK_019922, *cngc3* SALK_056832 (*cngc3-1*), *cngc3* SALK_066634 (*cngc3-2*), *cngc4* SALK_081369, *cngc5-1* SALK_149893, *cngc6-1* SALK_042207, *cngc9* GK_802D06 (*cngc9-1*), *cngc9* SAIL_736_D02 (*cngc9-2*), *cngc12-2* SALK_092622, *cngc17* SALK_041923). *bam1-3, clv1-20,* and *crn-3* have been described previously (B. J. DeYoung et al. 2006; Durbak and Tax 2011; Müller et al. 2008). *cngc9-1* GK-802-D06 R-GECO1 was a kind gift from Professor Petra Dietrich. The Ca^2+^ reporter line encoding the Ca^2+^ sensor R-GECO1 (Zhao et al. 2011) expressed under the control of the UBQ10 promotor (Norris et al. 1993) was a kind gift from Karin Schumacher (Keinath et al. 2015). *bam1-3, clv1-20, clv2-gabi* and *crn-3* were crossed to R-GECO1 and plants were identified by genotyping using the primers listed in supplementary table 1.

The promoter region of *CLE40* was amplified from Col-0 DNA and cloned into the *Promoter:H2B-Venus* destination vector by the use of the Gateway cloning system to yield *pCLE40:H2B-Venus*. The *pCLV1:CLV1-GFP* and *pBAM1:BAM1-GFP* plasmids were created using the Green gate cloning system (Lampropoulos et al. 2013). The entry plasmids (pGGB002, pGGE009, pGGF008) and the destination vector (pGGZ000) were used as provided from the manufacturer, whereas the promoter of *CLV1* and *BAM1* amplified from Col-0 DNA was cloned into pGGA000. The coding region of *CLV1* and *BAM1* amplified from Col-0 DNA was cloned into pGGC000. Primers used for generation of reporter lines are listed in supplementary table 1. Internal BsaI recognition sites were mutated by silent point mutations. Constructs were transformed into *Agrobacterium tumefacience* and the floral dip method (Clough and Bent 1998) was used to generate transgenic lines. *pCLE40:H2B-Venus* and *pCLV1:CLV1-GFP* were transformed into Col-0 while *pBAM1:BAM1-GFP* was transformed into the *bam1-3* mutant background. The *pCRN:mCherry-H2B*/*pCLV2:Venus-H2B* reporter line has been described previously (Somssich et al. 2016).

### Growth conditions

Seeds were surface sterilized with chlorine gas and incubated in 0.1 % (w/v) agarose for 2 days at 4 °C before plating onto 1/2 MS medium with 1 % (w/v) sucrose. Plates were incubated with constant light at 21 °C for 4-10 days. For root analyses, the plates were incubated vertically. For peptide treatment of roots plates containing a final concentration of 50 nM to 200 nM CLE40p were used and root length was analysed 10 dag.

### Confocal laser microscopy

Confocal laser microscopy was carried out on a Leica TCS SP8 STED 3X microscope using a 20x multi NA 0.75 objective, a Zeiss LSM 780 using a Plan-Apochromat 40x/1.0 DIC M27 objective and a Zeiss LSM 880 40x water / NA1.2 objective. Counterstaining of cell walls was used during image acquisition of reporter lines by staining roots in 2,5 µM propidium iodide for 5 min.

### Ca^2+^ imaging using the R-GECO1 sensor

[Ca^2+^]_cyt_ in roots were detected using plants expressing the red-emitting single-fluorescent protein R-GECO1 as a Ca^2+^ sensor (Keinath et al. 2015). Confocal laser scanning microscopy was performed on a Leica TCS SP8 STED 3X microscope using a 20x multi NA 0.75 objective and a Zeiss LSM880 Airyscan using a Plan-Apochromat 20x air NA 0.80 objective. The Ca^2+^ sensor R-GECO1 was excited with a white light laser at 561 nm and its emission was detected at 590 nm to 670 nm using a HyD detector (Leica) and an EMCCD iXon Ultra detector (Zeiss). Laser power was chosen for each experiment to maintain comparable intensity values and to avoid oversaturation. Images were recorded with a frame rate of 5 s at 600 Hz. Sample mounting was performed as previously described (Krebs and Schumacher 2013). Seedlings expressing the R-GECO1 Ca^2+^ sensor were grown on half strength Hoaglands or half strength MS agar for 10-11 days. Seedlings were incubated overnight in growth medium at 21 °C and continuous light conditions before the day of imaging. Peptides were prepared in two-fold concentration in half strength Hoaglands medium. The peptides were added in a 1:1 (final concentration 1 µM) volume ratio to the imaging chamber. ATP was prepared in a 100-fold concentration in 100 mM Tris pH 7.0 and added as a last treatment in a 1:100 volume ratio (final concentration 1 mM) to the imaging chamber. Image processing was performed in FIJI (Schindelin et al. 2012). These steps are: background subtraction, Gaussian blur, MultiStackReg v1.45 (http://bradbusse.net/sciencedownloads.html), 32-bit conversion, threshold. Royal was used as a look up table. Fluorescence intensities of indicated ROI s were obtained from the 32-bit images. Normalization was done using the following formula ∆F/F = (F-F_0_)/F_0_ where F_0_ represents the mean of at least 1 min measurement without any treatment. R-GECO1 experiments were performed independently in two different laboratories. The measurements gave identical results and all results were therefore combined in the data presentation. R-GECO1 measurements were performed at the Center for Advanced imaging (CAi) at HHU and at the NorMIC Imaging platform at the University of Oslo.

### Inhibitor treatment

1 M LaCl3 (Sigma-Aldrich) in water, 100 mM EGTA (Sigma-Aldrich) in water and 1M BAPTA (Sigma-Aldrich) in DMSO stock solutions were used at a final concentrations of 5 mM LaCl3, 100 µM EGTA and 2 mM BAPTA. Seedlings were preincubated in growth medium solution containing EGTA and BAPTA for 30 min, and LaCl3 for 2h before Ca^2+^ imaging was recorded.

### Peptide sequences

Peptides used in this study were ordered from Thermo Scientific. Peptide sequences are listed in supplementary Table 2.

### Movie

#### Movie 1

[Ca^2+^]_cyt_ dynamics in R-GECO1 expressing Col-0 wildtype root in response to 1 µM CLE40p. Roots 10 dag were treated with 1 µM CLE40p and changes in [Ca^2+^]_cyt_ was recorded over time. Response shown as normalized fluorescence intensities (∆F/F). Images were recorded with a frame rate of 5 s. Movie corresponds to measurements shown in Fig.1 A,B. Treatment with 1 mM eATP was used as a control, corresponds to Fig.1 C. Time shown as, hh:mm:ss. The first 5 min, roots were incubated in Hoagland’s medium and serve as background measurement.

#### Movie 2

[Ca^2+^]_cyt_ dynamics in R-GECO1 expressing Col-0 wildtype root in response to 1 µM CLE40p after preincubation in 100 µM EGTA for 35 minutes. Roots 11 dag were preincubated in EGTA and then transferred to the microscope. Roots were treated with 1 µM CLE40p and changes in [Ca^2+^]_cyt_ was recorded over time. Response shown as normalized fluorescence intensities (∆F/F). Images were recorded with a frame rate of 5 s. Movie corresponds to measurements shown in Fig.3 C. Treatment with 1 mM eATP was used as a control, corresponds to Fig.3 C. Time shown as, hh:mm:ss. The first 5 min, roots were incubated in Hoagland’s medium with 100 µM EGTA and serve as background measurement.

#### Movie 3

[Ca^2+^]_cyt_ dynamics in R-GECO1 expressing *clv2-gabi* root in response to 1 µM CLE40p. Roots 10 dag were treated with 1 µM CLE40p and changes in [Ca^2+^]_cyt_ was recorded over time. Response shown as normalized fluorescence intensities (∆F/F). Images were recorded with a frame rate of 5 s. Movie corresponds to measurements shown in supplemental Fig.2 C. Treatment with 1 mM eATP was used as a control, corresponds to supplemental Fig.2 C. Time shown as, hh:mm:ss. The first 5 min, roots were incubated in Hoagland’s medium and serve as background measurement.

## Supplementary Tables

**Supplementary Table 1:**
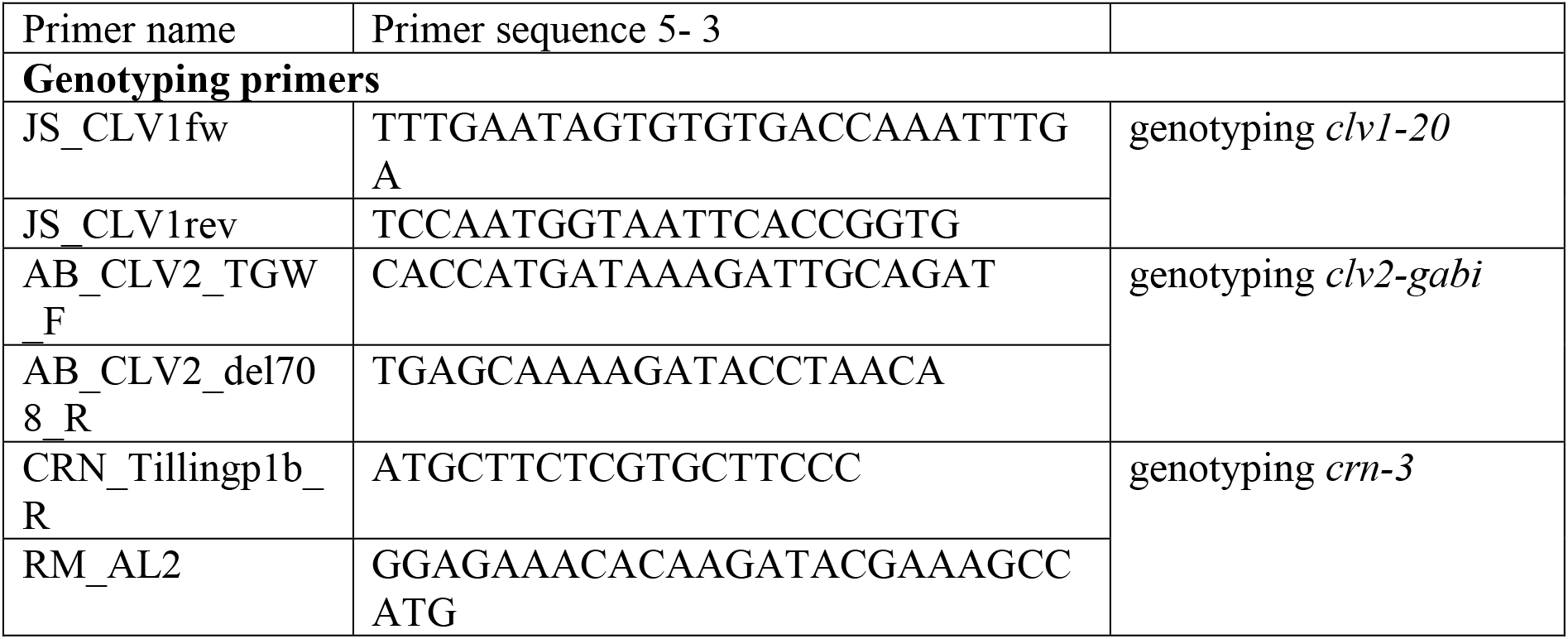
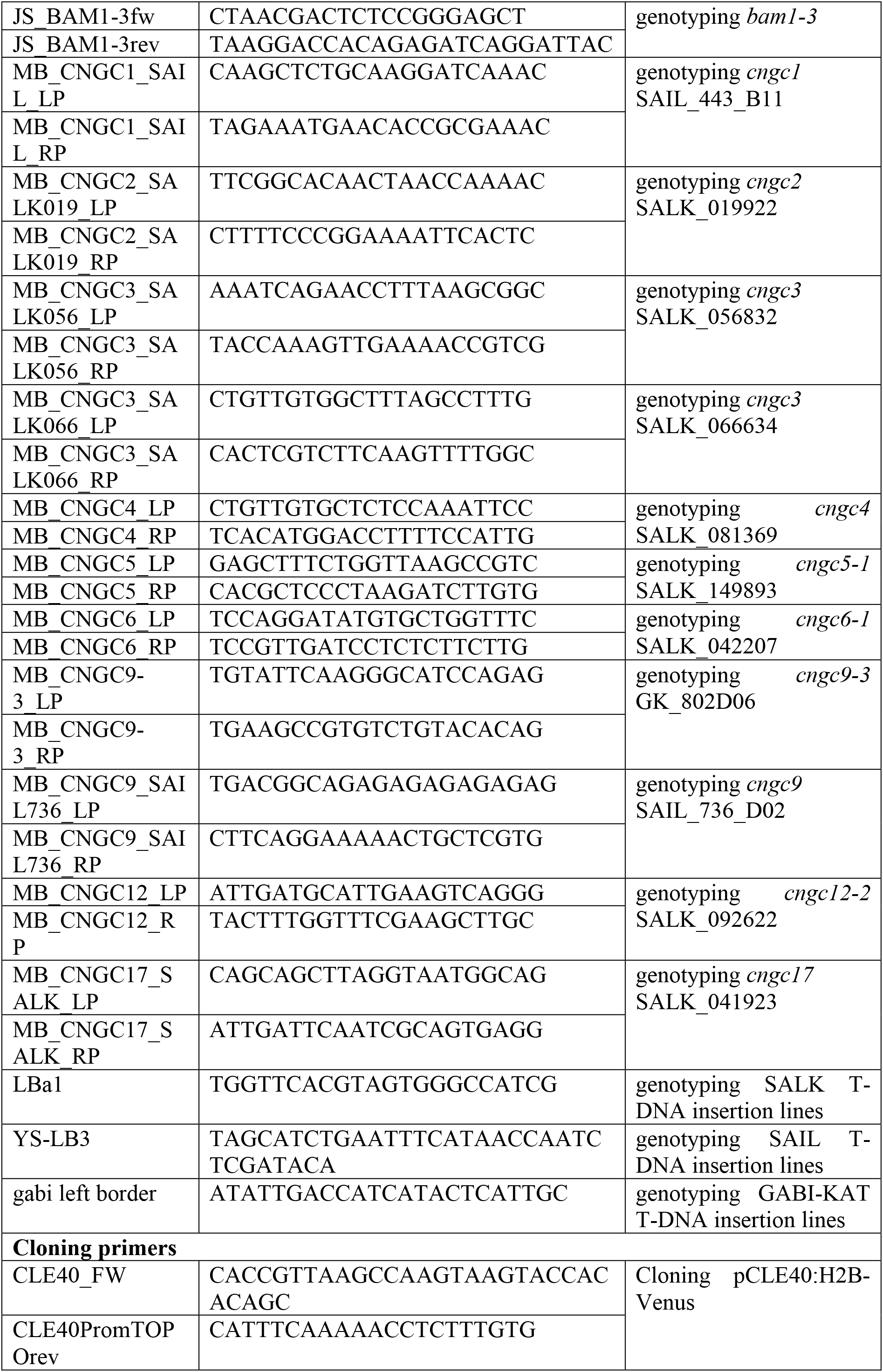
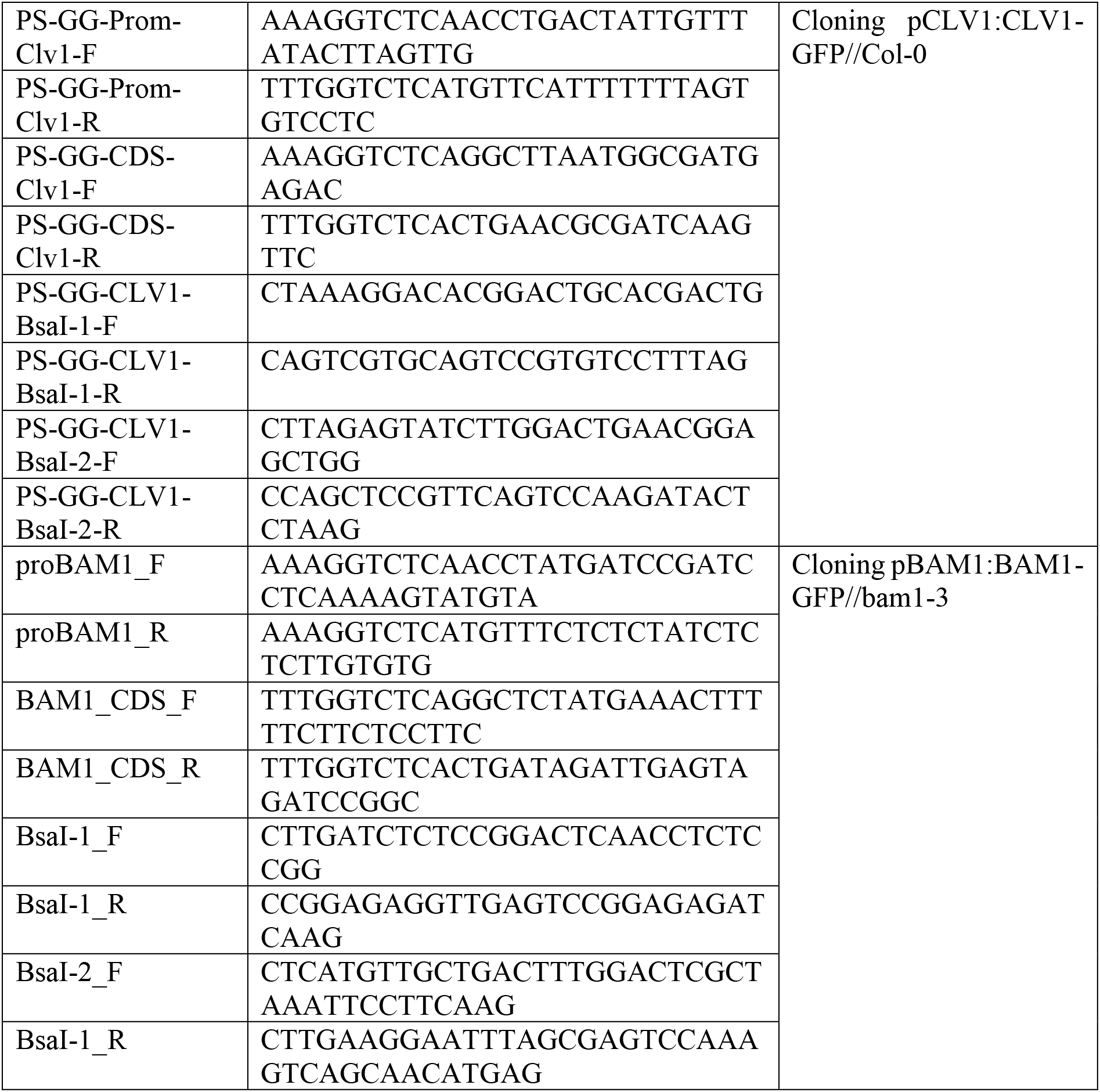
Primers used for genotyping and generation of reporter lines

**Supplementary Table 2:**
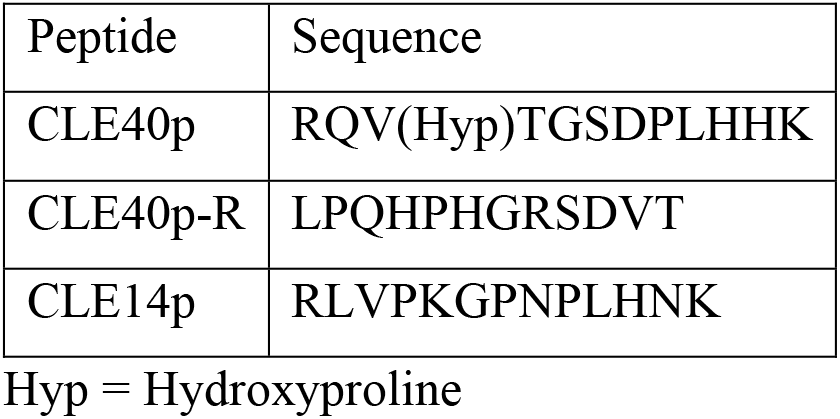
Peptide sequences.

## Acknowledgements

We thank Silke Winters and Carin Theres for CLE40p root assays and microscopy and Katharina Bursch for help with plant crosses to the R-GECO1 sensor. We are grateful to Dr. Marc Somssich for providing the *pCRN:mCherry-H2B* and *pCLV2:Venus-H2B* lines and to Professor Dr. Petra Dietrich for sharing *cngc9-1* R-GECO1 seeds with us. We thank Dr H. G. Nisse for critical reading of this manuscript. We thank the CAi Center for Advanced Imaging at HHU and the NorMic Imaging platform at UiO for the use and technical support. Work by R. Simon and M.B. was supported through CEPLAS (DFG, EXC2048). Work by V.O. and M.A.B. was supported by the Research Council of Norway (grant 230849).

## Author contributions

M.B. designed experiments, performed Ca^2+^ imaging, created R-GECO1 lines, performed CLE40p root assays, analysed Ca^2+^ imaging, genotyped mutant lines. V.O. performed Ca^2+^ imaging, analysed Ca^2+^ imaging, created R-GECO1 lines in *clv* mutant background. K G-P. performed CLE40p root assays. P.S. G.D. and J.S. performed gene expression studies. M.A.B. and R.S. designed experiments and drafted the manuscript. R.S., M.B. and M.A.B. wrote the paper with input from all authors.

## Supplementary figures

**Supplementary Fig.1:**
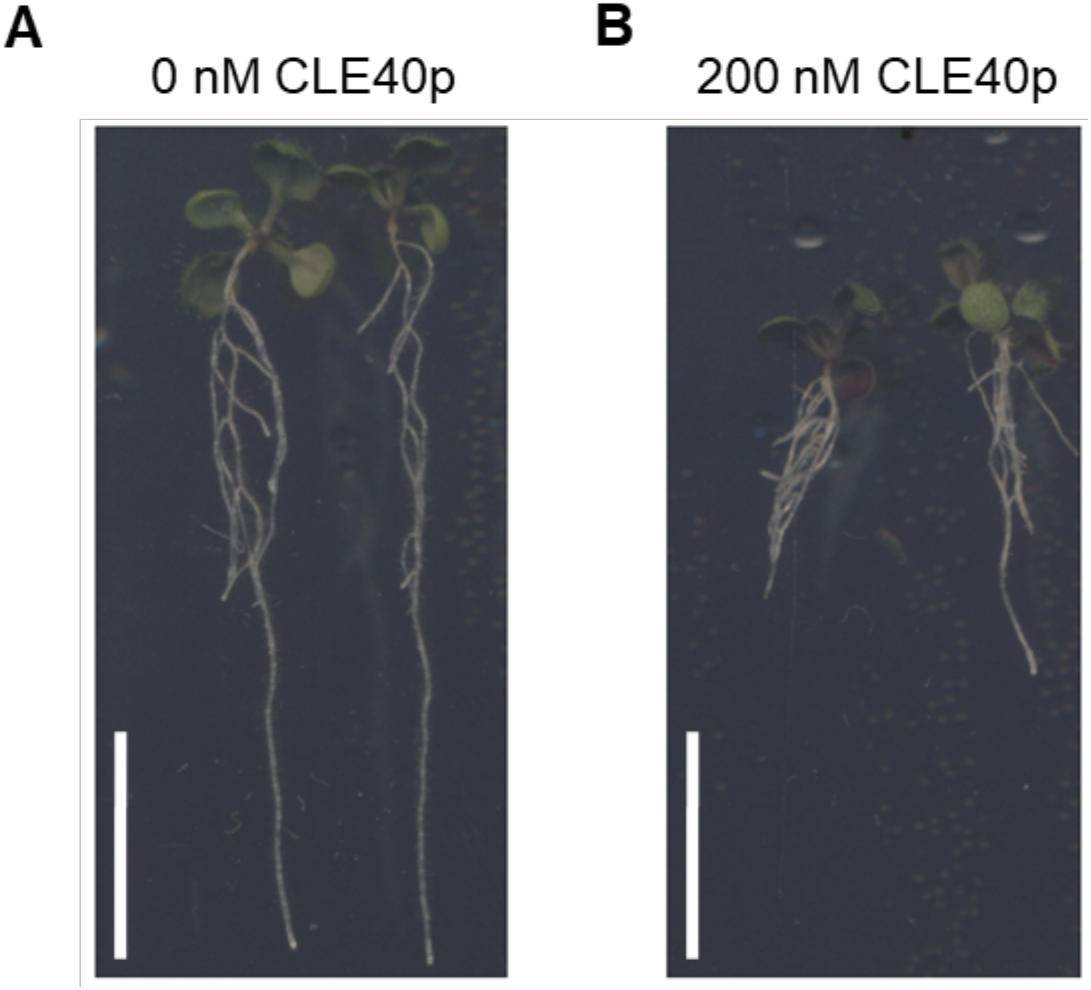
Phenotypes of 10 dag old WT (Col-0) seedlings grown on **A**, medium without CLE40p and **B**, medium with 200 nM CLE40p. Notice shorter roots in the presence of CLE40p. Scale bar = 1 cm

**Supplementary Fig. 2:**
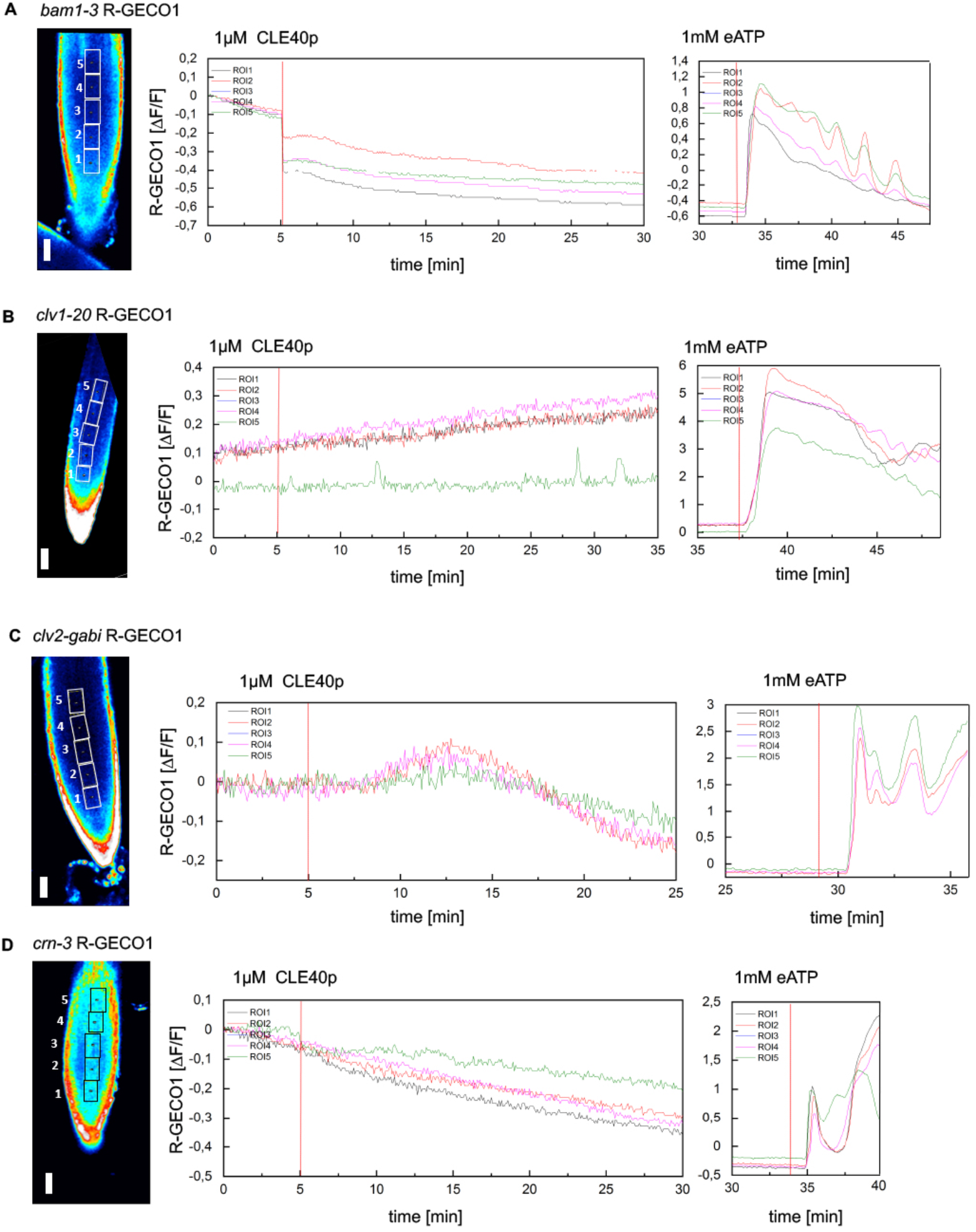
Representative response of the changes in [Ca^2+^]_cyt_ observed when roots expressing *R-GECO1* in the *bam1*, *clv1*, *clv2* and *crn* mutant backgrounds, respectively, are treated with 1 µM CLE40p followed by 1 mM eATP. Regions of interest (ROI) 1-5 (outlined in rectangles) were used to measure normalized R-GECO1 fluorescence intensities (∆F/F) and are represented in the graphs as dynamic responses over time. Red line at 5 min indicates application of CLE40p, followed by an additional red line indicating application of eATP, used as an internal positive control. No change in [Ca^2+^]_cyt_ response is observed in ROI 1-5 of: **A**, *bam1-3 RGECO1* roots (n = 13). **B**, *clv1-20 RGECO1* roots (n = 10). **C**, *clv2-gabi RGECO1* roots (n = 14) and **D**, *crn-3 RGECO1* roots (n = 14). Addition of eATP leads to increased fluorescence intensity representing an increase in [Ca^2+^]_cyt_ for all genotypes. scale bare = 50 µM

**Supplementary Fig. 3:**
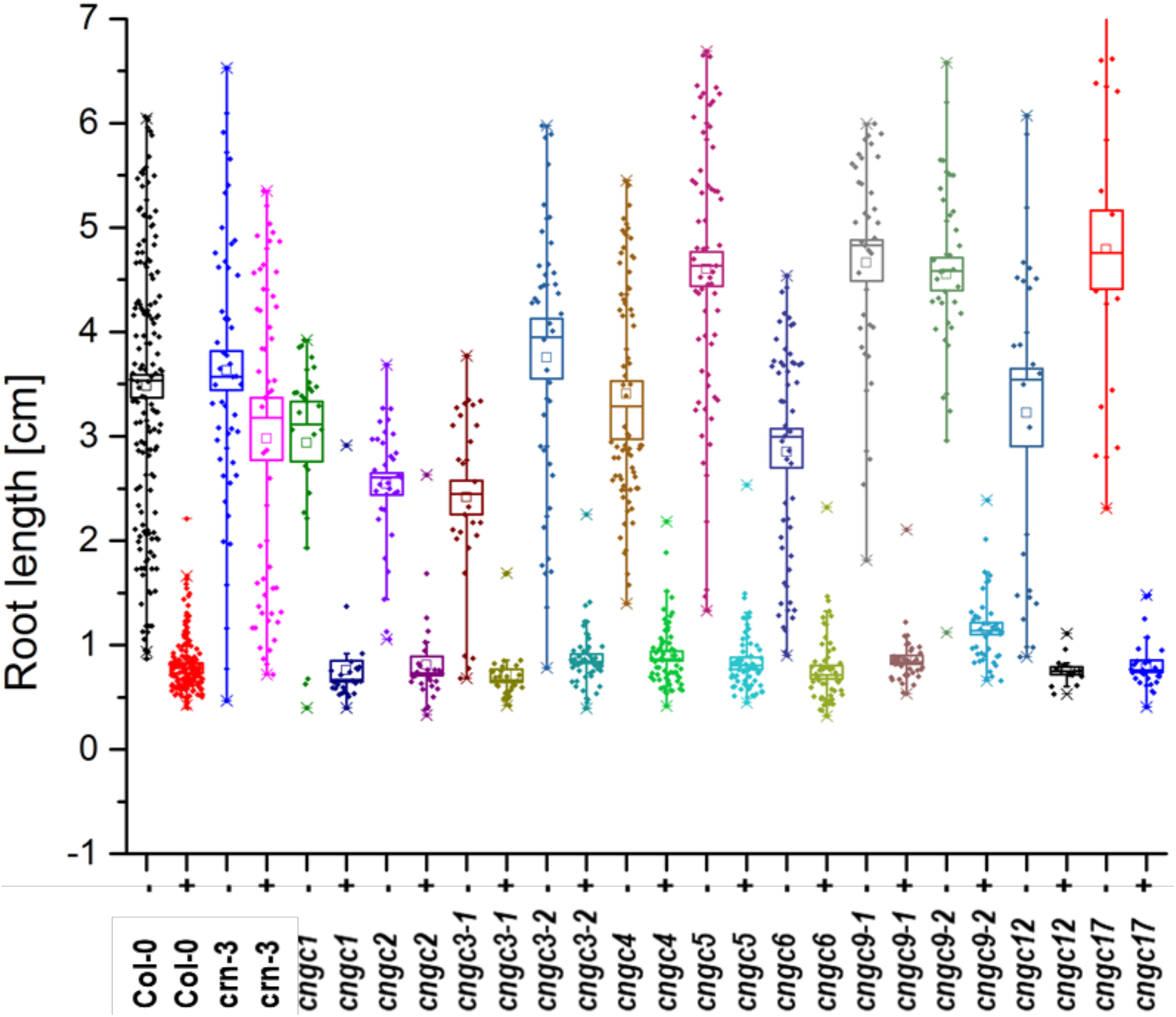
Impact of CLE40p treatment on different genotypes. Roots grown on medium with no peptide or on medium with a final concentration of 200 nM CLE40p were measured (in cm) 10 dag. All of the *cngc* mutants behaved like Col-0 WT and were responsive to CLE40p treatment showing a reduction in root length in the presence of CLE40p compared to untreated roots. The *crn-3* mutant was partially insensitive to CLE40p treatment.

